# Progressive neurodegeneration of the locus coeruleus drives rapid eye movement sleep fragmentation

**DOI:** 10.64898/2026.10.12.758491

**Authors:** Noam Zelinger, Noa Regev, Eviyatar Burian, Uddi Kimchy, Noam Gabay, Yuval Nir

## Abstract

Sleep disturbances are common early features of neurodegenerative disorders such as Alzheimer’s disease (AD) and Parkinson’s disease (PD). AD progression is also characterized by early and substantial degeneration of the noradrenergic locus coeruleus (LC), beginning with tau pathology and progressing to major neuronal loss. Although LC degeneration and sleep disruption progress in parallel, whether LC degeneration causally contributes to sleep abnormalities, and which sleep changes are associated with this process, remain unresolved. Human studies are largely correlative, while existing rodent models poorly reproduce progressive LC degeneration. Here, we employed progressive Caspase-3 (Casp3)-mediated LC ablation, with longitudinal sleep and calcium fiber photometry to monitor LC activity across defined stages of degeneration in adult female DBH-Cre mice. Histological analysis demonstrated graded, time-dependent LC degeneration, yielding mild degeneration (∼25% neuronal loss) by day 8 post Casp3 injection and moderate degeneration (∼50% loss) by day 18. Relative to baseline and to control mice, LC degeneration led to REM sleep fragmentation, emerging already at the mild degeneration stage. By the moderate stage, total REM sleep during light hours had significantly increased, while EEG theta power during REM sleep progressively declined across all stages of degeneration. Degeneration also blunted state-dependent LC calcium activity, significantly reducing the contrast between wakefulness and REM signal. Together, these results establish Casp3-mediated ablation as a model of progressive LC neurodegeneration and identify REM fragmentation and blunted state-dependent LC activity as its early functional signatures, providing a platform for studying how degeneration of neuromodulatory systems shapes sleep in a neurodegenerative disease-like state.

## Introduction

Neurodegenerative disorders such as Alzheimer’s Disease (AD) and Parkinson’s disease (PD) involve parallel sleep disruptions and degeneration of the monoaminergic neuromodulatory systems. In AD, cognitive decline and memory impairments constitute the primary clinical symptoms; however, sleep disturbances are also highly prevalent, affecting ∼60% of AD patients (compared to ∼15% in healthy aging)^1,2^ and are among the earliest detectable symptoms, often appearing years before memory deficits and clinical diagnosis^3,4^. Sleep assessment using polysomnography (PSG) shows progressive deterioration of several sleep features across the progression from healthy aging, through mild cognitive impairment (MCI), to advanced AD, which correlates with AD biomarkers^5^. Such sleep alterations include changes in macro-architecture involving reduced total sleep time, increased sleep fragmentation, changes in rapid eye movement (REM) and non-REM sleep, and in REM sleep latency^6^. Additional changes in micro-architecture include altered sleep spindles and slow waves that are associated with cognitive impairment ^7^.

Co-occurring with sleep disruptions, AD pathology also involves neurodegeneration of the Locus Coeruleus (LC)^8^, a fundamental sleep-wake regulator. The LC is a compact pontine nucleus that serves as the brain’s primary source of noradrenaline (NA) to promote arousal. In AD, the LC is the primary site for the accumulation of hyperphosphorylated tau protein (p-tau) before pathology is observed in the medial temporal lobe and episodic memory deficits appear^9^. Eventually, p-tau forms the neurofibrillary tangles (NFTs) that are a hallmark of the disease^10^. Progressive degeneration leads to an approximately 30% reduction in LC volume by the MCI or early AD stage^11–13^, eventually reaching a 60–80% neuronal loss in advanced AD^14^.

It has long been suspected that parallel sleep alterations and LC degeneration in AD may be related ^15^. Some posit that early LC dysfunction *causally* leads to sleep disturbances, which in turn can accelerate further neurodegeneration via a vicious cycle. However, other factors could drive LC and sleep changes in parallel, leading to concurrent changes without a causal relationship. Mechanistically understanding whether and how LC degeneration affects sleep remains elusive, as human research remains correlative and evidence relies on indirect measures such as postmortem data, magnetic resonance imaging, or pupillometry^16–18^.

Investigating AD-related LC-sleep relationships in rodents could potentially elucidate causality, but challenges remain. LC Progressive degeneration is not replicated by transgenic AD mouse models ^19^ or by lesion studies that cause abrupt and excessive LC damage, which have yielded inconsistent results. For instance, LC lesions by 6-Hydroxydopamine (6-OHDA) produce either no effect on sleep stages and electroencephalogram (EEG) properties^20,21^, transient reduction in wakefulness^22^, or long-term increase in REM sleep^23^. Similarly, the neurotoxin N-(2-chloroethyl)-N-ethyl-2-bromobenzylamine (DSP-4), which targets noradrenergic projections, leads to short-lived REM suppression^24,25^, with either circadian-dependent fragmentation^26^ or no changes at all to sleep architecture^27^.

We sought to bridge the gap between acute experimental animal models of LC damage and progressive LC degeneration in human pathology by experimentally inducing progressive LC degeneration and testing the possible causal effects on sleep. To this end, we utilized a Cre-dependent Caspase-3 (Casp3) viral approach in DBH-Cre mice to induce apoptosis (programmed cell death) specifically in the LC. We induced a programmed progressive partial ablation of the LC to better approximate the gradual neurodegenerative timeline observed in patients. By implementing longitudinal sleep polysomnographic monitoring (via EEG, electromyogram (EMG), and video) and LC activity monitoring (via fiber photometry of bulk calcium activity), we tracked sleep architecture and LC activity in the same animals across multiple time points. Our approach successfully led to gradual lesioning of the LC, with around 25% cell loss after one week and 50% cell loss after two weeks. Such structural damage was associated with sleep abnormalities, particularly REM sleep fragmentation and theta power reduction, and with blunted differences in LC calcium activity across vigilance states.

## Methods

All experimental procedures, including animal handling, surgery, and experiments, followed the NIH Guide for the Care and Use of Laboratory Animals and were approved by the Institutional Animal Care and Use Committee (IACUC) of Tel Aviv University (approval TAU-MD-IL-2303-118-4). Adult 14-20 week-old DBH-cre female mice (B6.FVB(Cg)-Tg(Dbh-cre)KH212Gsat/Mmucd, 036778-UCD) were used. Mouse strain was obtained from the Mutant Mouse Resource and Research Center (MMRRC) at University of California at Davis, an NIH-funded strain repository, and was donated to the MMRRC by MMRRC at University of California, Davis. Made from the original strain (MMRRC:032081) donated by Nathaniel Heintz, Ph.D., The Rockefeller University, GENSAT and Charles Gerfen, Ph.D., National Institutes of Health, National Institute of Mental Health. Mice were housed in transparent Plexiglas cages under controlled conditions: temperature (20°-23°C), humidity (40-70%), and a 12-hour light/dark cycle (lights on at 10:00 AM). Food and water were provided ad libitum.

### Experimental Design

This study comprised two experimental phases.

#### Phase 1 (histology-only)

In this phase, two separate cohorts were used to characterize Casp3-mediated LC degeneration by employing unilateral virus injection and comparison to the contralateral hemisphere (as seen in **Fig. 1**): a virus dilution verification cohort (phase 1.1) was used to establish the relationship between viral titer and lesion severity, and a timeline characterization cohort (phase 1.2) was used to map the progressive time course of degeneration following a fixed viral dose. Based on this characterization, we selected a dilution and injection volume expected to produce partial, graded LC degeneration suitable for subsequent functional studies.

**Figure 1.**
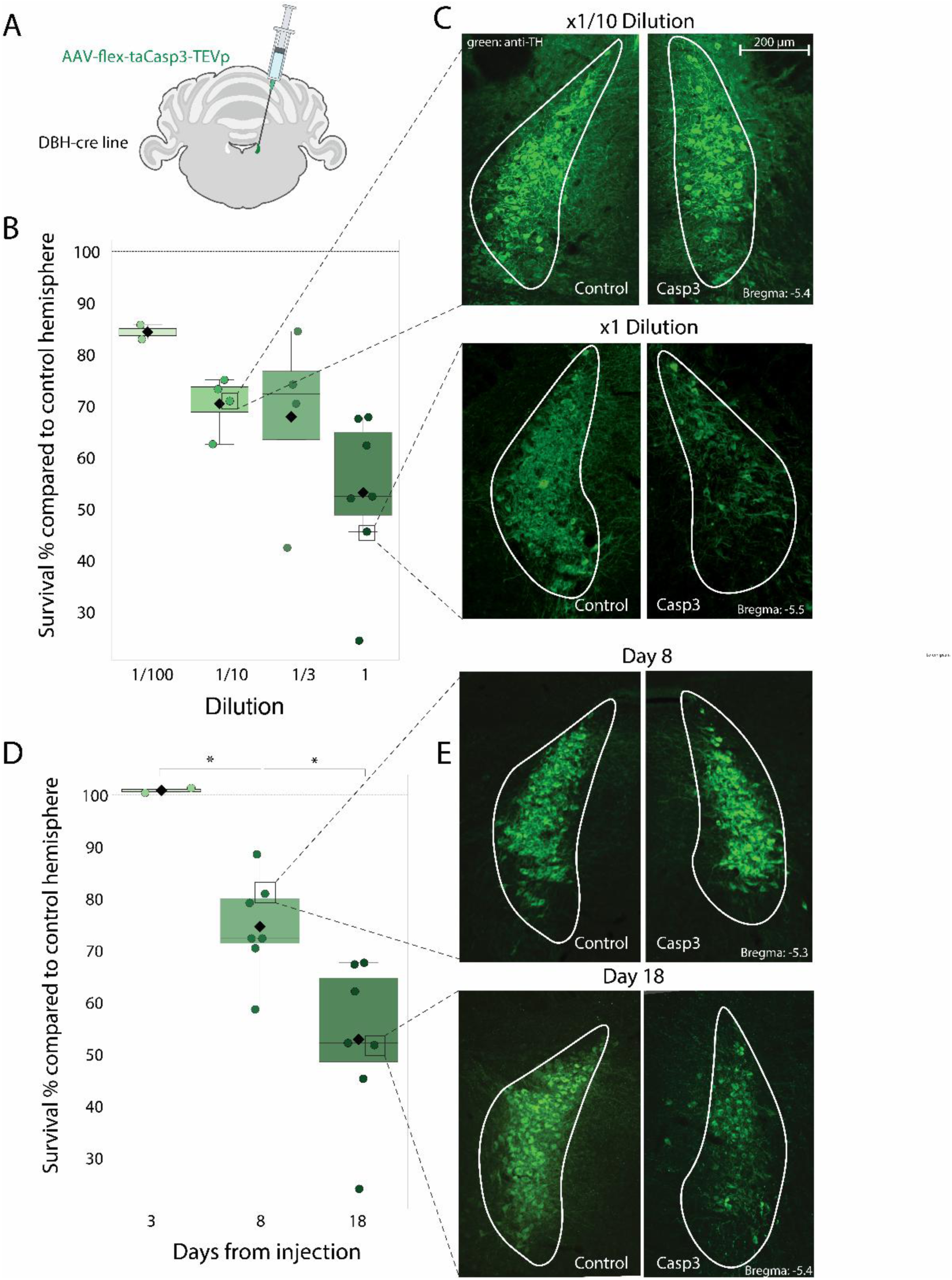
Characterization of Casp3-mediated progressive LC degeneration. (A) Schematic of unilateral Casp3 injection into the LC of DBH-Cre mice. (B) LC survival (y-axis) across viral dilutions (x-axis) at 14–21 days post-injection reveals a dose-response relationship of increased damage with higher virus concentration (F = 3.31, p = 0.05). (C) Representative brain sections with anti-TH staining for 1:10 dilution (top) and undiluted 1:1 (bottom) concentrations. Scale bar = 200 µm. (D) LC survival (y-axis) across different durations from undiluted Casp3 injection (x-axis) reveals a dose-response relationship of increased damage with longer elapsed time (one-way ANOVA, F = 13.60, p = 0.0006; Welch’s t-test, Day 3 vs. 8: p = 0.0003; Day 8 vs. 18: p = 0.0095). Horizontal lines and error bars mark mean ± SEM; circles mark values in individual mice (Supplementary Tables 1 and 2). (E) Representative anti-TH staining for mild LC damage at day 8 (top) and moderate LC damage at day 18 (bottom).

#### Phase 2 (sleep-photometry experiment)

A separate third cohort of Casp3-injected and control mice underwent bilateral LC injections together with EEG, EMG, and fiber photometry implantation (as seen in **Fig. 2A**), allowing sleep architecture and LC activity to be tracked within the same animals across defined stages of degeneration. Twenty-one days before the start of the experiment, GCaMP was injected to permit expression. On day 0, we implanted EEG, EMG, and photometry cannulas and injected Casp3 virus bilaterally (surgical procedures described below). Recordings started before neuronal damage to establish a within-subject baseline. Animals were allowed one day of recovery in their home cage, followed by one habituation day in a sound-attenuated chamber (–55 dB, H.N.A.) while tethered to the EEG/EMG cable and fiber-optic patch cord. Electrophysiology and photometry data were recorded for 22 hours, beginning 30 minutes after light onset (10:30 AM). These 22-hour recording sessions were performed on Days 3, 8, and 18 post-injection to capture progressive stages of LC degeneration (baseline, mild, and moderate, respectively; day 3 was considered baseline since we didn’t observe any degeneration 3 days after casp-3 injection). Following the final recording session, animals were euthanized for histological analysis. A total of 13 animals (experimental group: n=7; control group: n=6) were used in this experiment.

**Figure 2.**
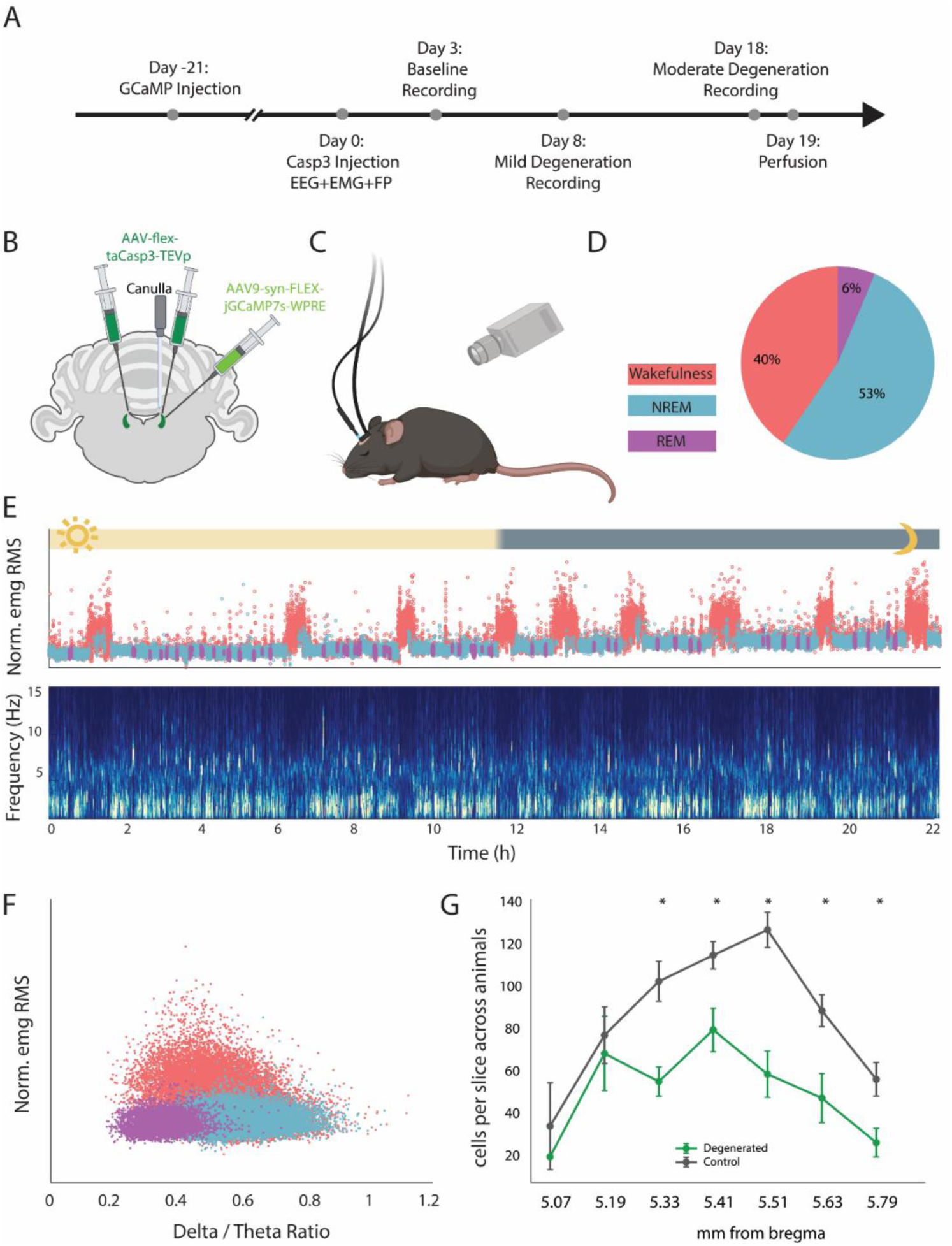
Experimental design. (A) Timeline of the sleep experiment. (B) Schematic of bilateral LC injections for Casp3-mediated degeneration and GCaMP-based fiber photometry. (C) Recording setup in a freely behaving mouse including combined EEG, EMG, fiber photometry, and video monitoring. (D) Proportion of time in each vigilance state during a representative recording. (E) Representative hypnogram (time-course of sleep/wake vigilance states) and EEG spectrogram (time-frequency decomposition) across a full recording session. Note epochs of NREM sleep (cyan) enriched with slow wave activity, interleaved with wakefulness (red, high EMG RMS levels) and sporadic intervals of REM sleep (purple). (F) Scatter plot of EMG RMS (y-axis) and EEG delta/theta power ratio (x-axis) reveals separation of vigilance states. (G) Average of the animals’ mean number of cells in a slice across the Dorsal-Ventral axis (Welch’s t-tests, BH-FDR correction for multiple comparisons).

### Surgeries

Before all surgical procedures, mice were anesthetized with isoflurane (3% for induction, 1.2% for maintenance) and received lidocaine (3mg/kg) locally and buprenorphine (0.05mg/kg) and carprofen (5mg/kg) systemically for analgesia. They were placed in a stereotaxic frame and craniotomy was performed, under microscopic control using a high-speed surgical drill, for virus injections and optic fiber/EEG screw implantations. Virus injections were delivered using a UMP3 Microsyringe Injector and Micro4 Controller pump with 33-gauge NanoFil needles (World Precision Instruments, USA) at a rate of 80 nl/min. The LC was targeted using the following coordinates relative to bregma, based on Allen Brain Atlas: antero-posterior (A/P): -5.4 mm; medio-lateral (M/L): 1 mm; dorso-ventral (D/V): - 3.3 mm and -3.5 mm relative to the brain surface.

#### Casp3 virus injections for histology-only experiments (Phase 1)

The Cre-dependent Casp3 virus (pssAAV-2-hEF1α-dlox-(pro)taCasp3_2A_TEVp(rev)-dlox-WPRE-hGHp(A); ETH, Zurich, Switzerland) was injected unilaterally at the coordinates described above. At the end of surgery, the scalp was sutured, and mice were returned to their home cages. Our target viral load was 80 nL of undiluted stock (titer 10¹² vg/mL). To improve volumetric reliability while preserving this target load, all injections in this study were delivered as a 320 nL volume of fourfold-diluted virus, rather than 80 nL of undiluted stock. For the virus dilution verification cohort (phase 1.1), we conducted additional experiments across a range of viral dilutions (1:99, 1:9, and 1:2 relative to stock, referred to as x1/100, x1/10, and x1/3 in Results and figures) to characterize the relationship between viral concentration and lesion extent (n=17 female mice). For the timeline characterization cohort (phase 1.2), mice were injected unilaterally with a standard fourfold dilution (original stock titer, 320 nL total volume) and euthanized at 3, 8, and 18 days post-injection to characterize the progressive cellular degradation induced by the virus (n=16 female mice).

#### GCaMP virus injections (Phase 2)

For sleep–photometry experiments, in a first surgical procedure performed 21 days before experiments (Fig. 2A), cre-dependent GCaMP (AAV9.syn.FLEX.jGCaMP7s.WPRE) was injected unilaterally into the LC at a total volume of 250 nL. Following injection, the scalp was sutured, and mice were returned to their home cages. For the control group, WT littermates underwent identical viral injections.

#### EEG, EMG, and optic fiber preparation (Phase 2)

In the second surgical procedure performed on Day 0 (Fig. 2A), two screws (1 mm diameter) were implanted over the left hemisphere for EEG recording: one over the parietal plate and another above the cerebellum/posterior parietal lobe used as a combined reference and ground. EMG signals were recorded via two single-stranded stainless-steel wires inserted bilaterally into the neck muscles in a bipolar reference configuration. EEG and EMG wires were soldered onto a custom headstage connector. Subsequently, an optic fiber Cannula (MFC_200/240-0.22_5mm_MF1.25_FLT, Doric Lenses) was implanted above the LC, and C&B Metabond dental cement was applied to secure all screws, wires, and the Cannula to the skull. In the same surgery, the Cre-dependent Casp3 virus was injected bilaterally into the LC at the same coordinates as previously described.

### Sleep polysomnography via EEG, EMG, and video recordings (Phase 2)

EEG and EMG were digitally sampled at 1,017.25 Hz (1700 Differential AC Amplifier, A-M Systems) and notch-filtered at 50Hz and 100Hz to remove line noise and harmonics. Simultaneous video data (for sleep and free behavioral assessments) were captured with a USB webcam and synchronized with electrophysiological and photometric data.

### Fiber photometry recordings (Phase 2)

Fiber photometry data were collected as previously described in Atlan et al. 2024^28^ using a single-site system (Doric Lenses, Canada) adapted to two excitation LEDs at 465nm (GcaMP signal) and 405nm (isosbestic control channel). The two channels were monitored simultaneously by connecting the LEDs to a 4-port minicube (FMC4, Doric Lenses; containing dichroic mirrors and cleanup filters matched to excitation and emission spectra) via an attenuating patch cord (400 µm core, NA=0.37-0.48). LEDs were controlled by drivers that sinusoidally modulated 465 nm and 405 nm excitation at 217 Hz and 330 Hz, respectively, enabling lock-in demodulation. Zirconia sleeves were (MFP_400/430/1100-0.57_1m_FC-ZF1.25_LAF, Doric Lenses) attached the fiber-optic patch cord to the animal’s implanted cannula. Data were sampled at 1,017.25 Hz and processed using an RZ2 BioAmp Processor unit and collected using Synapse software (Tucker-Davis Technologies). To achieve 22-hour recordings while reducing adverse effects of prolonged LED activation (e.g., phototoxicity and photobleaching), we employed intermittent fiber photometry with alternating 1-hour periods of LEDs on and LEDs off, as previously described in Matosevich et al. 2025^29^. Of the seven Casp3-injected animals, three were excluded from this analysis: one lacked a usable isosbestic channel in the baseline recording and could not be normalized as in other sessions; one showed no anti-correlation between the photometry signal and sigma power, indicating the absence of a reliable LC GCaMP signal; and one showed a slow within-hour signal drift, consistent with residual photobleaching, which confounded comparisons between vigilance states. Data from the remaining four animals were included, representing reliable and state-differentiated signal.

### Histology

Under very deep isoflurane anesthesia (4%) combined with a ketamine-xylazine dose (ketamine 87 mg/kg, xylazine 13 mg/kg), mice were perfused intracardially with saline (0.9% NaCl, 1 ml/g) followed by 4% PFA(Merck). Brains were then extracted and fixed for 48 h in 4% PFA at 4°C. Coronal brain sections encompassing the LC were cut using a vibrating microtome (Leica VT1000 S). For the Sleep-Photometry Experiment and Virus Dilution Verification, sections were cut at 60 µm and stored free-floating in PBS. For the Timeline Characterization cohort, tissue was sectioned at 20 µm to accommodate automated slide-scanner imaging.

### Immunohistochemistry

Viral expression and LC integrity were evaluated by immunostaining for tyrosine hydroxylase (TH). Sections were washed three times in PBS (Hylabs), permeabilized in PBST (PBS containing 0.1% Triton X-100; Merck), and blocked in PBST containing 20% normal goat serum (NGS; Vector Laboratories) for 1 hour at room temperature. Sections were incubated with primary antibody (chicken anti-TH, 1:300; ab76442, Abcam) in PBST with 2% NGS for 30 minutes at room temperature, followed by two overnights at 4°C. After three PBST washes, sections were incubated with secondary fluorophore-conjugated antibodies (AF488 goat anti-chicken, 1:500, ab150173, Abcam; or CF594 anti-chicken IgY, 1:500, SAB4600102, Merck) in PBST with 2% NGS for 1 hour at room temperature. Sections were washed three times in PBST and once in PBS, mounted onto glass slides, and coverslipped with fluorescent mounting medium containing DAPI (Fluoroshield™, F6057, Merck).

### Fluorescence Microscopy

For the Virus Dilution Verification cohort, image acquisition was performed using a 3i Marianas Spinning Disk Confocal System (Intelligent Imaging Innovations) to enable precise cell quantification. Z-stack images spanning 60 µm sections were acquired with a 20× air objective (0.4 NA) to capture LC cells across the Z-plane. For the Timeline Characterization and Sleep-Photometry Experiment cohorts, images were acquired using a VS200 Slide Scanner (Evident Scientific) with a 20× objective. Automated whole-slice scanning with image stitching was employed to ensure complete detection of remaining LC neurons without manual localization bias. Images were processed and analyzed using ImageJ^30^ and Cellpose^31^ software.

### Image-based cell quantification

For the Virus Dilution Verification and Timeline Characterization cohorts, unilateral injections were performed, allowing LC survival to be calculated relative to the intact contralateral LC. Fluorescent TH-positive LC cells were counted on both the injected and control sides using CellPose software with blinded manual verification. Total cell counts per side were aggregated for each mouse, and the percentage of surviving cells was calculated as: 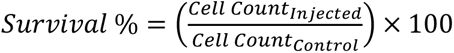. Mice exhibiting less than a 5% difference between control and injected sides were classified as injection failures and excluded from analysis (2 animals out of 18).

For the Sleep–Photometry Experiment cohort, bilateral injections were performed and internal contralateral control was not available. Instead, a blinded experimenter matched each coronal slice to its corresponding neuroanatomical level in the Allen Brain Atlas. Cell counts from bilateral Casp3-injected mice were compared against the WT control injected with Casp3, and against the non-injected hemisphere of all animals in the timeline characterization.

### Sleep data and scoring (Phase 2)

Continuous 22-hour EEG and EMG recordings were analyzed using a semi-automatic sleep-scoring method as described in ^32^, adapted from ^33^. For each animal, four non-consecutive hours from one session were manually scored, followed by manual scoring of a 1-hour segment for the remaining sessions. A convolutional neural net was trained using each animal’s manually scored sleep vector (tagged at a 1-second resolution), using the parietal EEG signal (band-pass filtered 0.3-30 Hz) and the EMG signal (band-pass filtered 10-50 Hz). The entire dataset was then classified into vigilance states (wake, NREM, and REM). The automatic classification results were visually inspected to ensure accuracy. Sessions with excessive artifacts or low signal-to-noise ratios (SNR) not suitable for automatic scoring (∼16% of the data) were manually scored for the full 22-hour duration.

### Fiber photometry data processing (Phase 2)

Data were detrended and normalized during 1-hour LED-ON periods following the method described by^34^. Briefly, the isosbestic channel was fitted to the GCaMP trace using a linear polynomial fit and subsequently used as the f_0_ reference point: 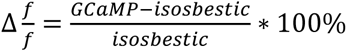. In addition, a degree-3 polynomial was fit to the normalized ΔF/F signal and subtracted to remove any slow trends that could represent residual signal bleaching. Mean fluorescence values for each vigilance state include only epochs longer than 5 seconds, with each 5-second segment averaged into a single value (e.g., a 30-second wakefulness epoch resulted in six mean values). To allow inter-animal comparisons, fluorescence signals were z-scored within each animal according to the baseline session’s mean and std, preserving within-animal relationships while eliminating between-animal variability.

### Statistical analysis

All statistical tests were performed in Python (SciPy v1.15.1, Pingouin v0.5.5, statsmodels v0.14.6). Significance was set at α = 0.05 throughout. Non-parametric tests were used to verify robustness without assuming normality. Where a parametric test was used, we used Welch’s t-test rather than Student’s t-test without assuming equal variances between groups.

LC degeneration time course and dilution effects were assessed with one-way ANOVA, followed by planned pairwise comparisons between consecutive stages using Welch’s t-test. In the sleep– photometry cohort, Casp3-injected animals were compared to WT littermates using Welch’s t-tests on per-animal mean LC cell counts, both overall and section-wise along the antero-posterior axis (Benjamini–Hochberg FDR correction^35^ across sections).

Within-group and between-group changes in sleep architecture and spectral power were tested with Wilcoxon signed-rank and Mann–Whitney U tests, respectively, with Bonferroni correction for multiple post-baseline comparisons per animal. Monotonic trends in REM theta and NREM delta power across baseline, mild, and moderate stages were assessed with Page’s L trend test for repeated measures. State-dependent LC photometry was analyzed with an exact permutation test on the Session × State interaction: for each animal, the change in mean Z-scored signal (moderate − baseline) was computed for Wake, NREM, and REM, and the assignment of these difference scores to states was permuted within animals to generate the null distribution for a Wake–REM contrast statistic (weights −1, 0, +1). As a converging analysis, we also report the Session × State interaction from a two-way repeated-measures ANOVA.

## Results

### Graded LC degeneration via Cre-dependent Caspase-3 ablation

To model progressive LC loss, Cre-dependent Caspase-3 (AAV-flex-taCasp3-TEVp) was injected unilaterally into DBH-Cre mice, using the intact contralateral LC as a within-subject control (**Fig. 1A**). We induced varying degrees of LC cellular loss by applying four degrees of dilution (1/100, 1/10, 1/3, and 1, corresponding to titers of 3.33*10^11^, 10^11^, 10^10^, and 10^12^ vg/mL respectively; volume 80nL). These parameters revealed a dose-response relationship between Casp3 and LC damage (**Fig. 1B, F** = 3.31, p = 0.05 via one-way ANOVA). Corresponding representative histological quantifications from individual mice show that, on average, surviving LC cells comprised 84.3% ± 1.4, 70.4% ± 2.8, and 67.8% ± 9.0 of the number of cells in the contralateral undamaged LC for dilutions of 1/100, 1/10, and 1/3, respectively (mean ± SEM, see **Supplementary Table 1** for details).

In the next phase, we continued to investigate the longitudinal dynamics of degeneration, focusing on the full-strength undiluted virus associated with substantial moderate-to-severe cell loss (53.1% ± 5.8, **Fig. 1C bottom**). Mice were sacrificed on days 3, 8, and 18 after Casp3 injection, and the extent of LC damage was quantified within each animal relative to the healthy contralateral LC. We found that elapsed time significantly increased the extent of LC damage (F = 13.60, p = 0.0006 via one-way ANOVA; **Fig. 1D**). While LC cell counts at day 3 were indistinguishable from the healthy contralateral LC (100.9% ± 0.4%, n=2), by day 8 they significantly dropped to 74.7% ± 3.6 (n=5; Welch’s t-test, t = 7.28, p = 0.0003). Neuronal degeneration progressed significantly further by day 18, reaching 53% ± 5.8 (n=5; Welch’s t-test, t = 3.20, p = 0.0095 vs. Day 8; **Fig. 1D, E**). These results establish a multi-stage Casp3 injection model of progressive LC loss at days 3, day 8, and day 18 that we will refer to as ‘baseline’, ‘mild degeneration’, and ‘moderate degeneration’ throughout the remainder of the manuscript. Next, we investigated how these three conditions may modulate sleep and LC-NE calcium activity.

### LC degeneration selectively disrupts REM sleep architecture and theta power

To assess how progressive LC degeneration affects sleep, mice received bilateral Casp3 injections and were monitored via polysomnography (EEG, EMG, and video) and LC fiber photometry. Sleep was scored according to established criteria, yielding labels of wakefulness, NREM sleep, REM sleep, and artifacts for each 1-sec interval (Methods).

We continuously recorded 22h of sleep/wake dynamics in the home cage at three temporal stages (corresponding to the degeneration model established above): at baseline (day 3) before degeneration, a recording at day 8 representing mild degeneration, and a recording at day 18 representing moderate degeneration (**Fig. 2A–C**).

**Fig. 2D–F** illustrates a representative distribution of time spent across vigilance states at baseline with 53%, 40%, and 6% of total time spent in NREM sleep, wakefulness, and REM sleep (1% of data contained artifacts), consistent with expected patterns^36^. LC cell count was significantly lower in Casp3-injected animals compared to controls, with experimental animals showing only 58% of control LC cell density on average across all sections (Welch’s t-test, t = –4.48, p = 0.0027). This reduction was consistent across most of the Anterior-Posterior axis, reaching significance at all positions except the two most anterior sections (**Fig. 2G**).

Next, we examined the effects of LC degeneration on sleep. We observed an array of changes specific to REM sleep as follows. During light hours, when mice are predominantly asleep, REM sleep was selectively disrupted by LC degeneration. We found that the median REM epoch length shortened significantly by 39.2% from baseline to both mild and moderate degeneration, with a concomitant increase of 41.5% in the number of REM epochs at both stages (Wilcoxon signed-rank test, all p = 0.031; **Fig. 3A, B**). Together, these effects represent fragmentation of REM sleep with more shorter epochs, and both effects were specific to the degeneration group, differing significantly from the corresponding change in control animals at both stages (Mann-Whitney U, all p < 0.03). By day 18 (moderate degeneration), total REM duration also increased significantly from baseline (p = 0.031; **Fig. 3C**); this change was relatively specific to the Casp3 group, trending towards statistical significance (p= 0.15) in the direct between-group comparison. During the dark hours, REM median epoch length was reduced under mild degeneration (p = 0.031; **Fig. S1A**), but no other metric showed a change specific to the degeneration group. Importantly, progressive LC ablation was rarely associated with robust changes in wakefulness or NREM sleep metrics compared to baseline (**Fig. S1B, D**) and even when we did observe such changes across time, they were not significantly different than those observed in control animals (Mann-Whitney U, p = 0.64 for each; **Fig. S1C, E**), indicating it was unrelated to the induced LC cellular damage.

**Figure 3.**
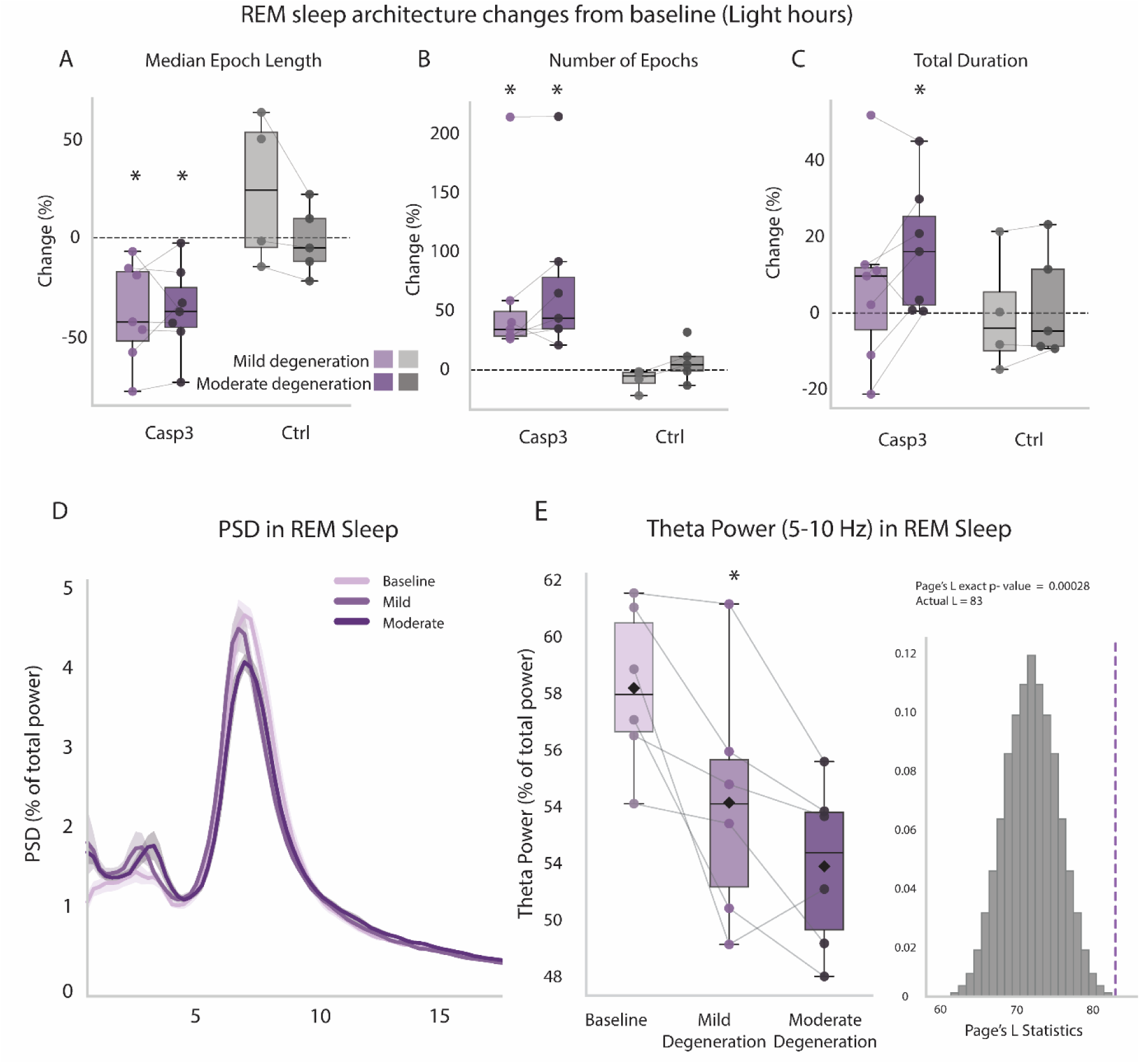
LC degeneration selectively disrupts REM sleep architecture and theta power during light hours. (A) Change in median REM sleep epoch length (y-axis, expressed as percent change from baseline) in experimental Casp3 mice (purple, left bars) versus control mice (gray, right bars) during the light phase when mice predominantly sleep. Lighter and darker colors mark mild and moderate degeneration, respectively. Note robust degeneration-induced decrease of REM epoch length. (B) Same as A for number of REM sleep epochs. (C) Same as A for the total duration of REM sleep. Asterisks indicate significant within-group change from baseline (Wilcoxon signed-rank, p_adj < 0.05. (D) Power spectral density (PSD) of the EEG during REM sleep. Colors as above. Note decrease in theta (5-10Hz) power with LC degeneration. (E) REM theta power across degeneration stages, showing a significant monotonic decrease (Page’s L trend test, L = 83, n = 6, p < 0.001). Data are mean ± SEM across mice.

Given the association between LC damage and REM sleep fragmentation, we hypothesized that REM sleep may be impaired in experimental mice beyond the gross metrics of epoch length and number. We therefore analyzed the power spectrum of EEG during REM sleep. The results revealed a significant monotonic decrease in theta (5–10 Hz) power across baseline, mild, and moderate degeneration (Page’s L trend test, L = 83, n = 6, p < 0.001; **Fig. 3D, E**), while the control group didn’t show a similar statistically significant trend (L = 52, n =4, p = 0.11). No corresponding trend was observed for NREM slow-wave power (0.5–4 Hz) across the same time points (L = 75, n = 6, p = 0.24; **Fig. S1F, G**).

### Cellular degeneration blunts state-dependent LC activity

To assess how structural LC degeneration affects the dynamics of LC activity across vigilance states, we examined LC population calcium activity using fiber photometry, focusing on differences between baseline and moderate degeneration (n=4, Methods).

Representative traces showed that LC dynamics showed multiple expected characteristics attesting to adequate measurements during long monitoring periods: elevated ΔF/F levels during wakefulness, variable levels during NREM sleep following infraslow (0.02Hz) oscillations where peaks were related to increased micro arousal propensity, and a sharp decline and flattening during REM sleep (**Fig. 4A**). Furthermore, during NREM sleep we observed the well-documented^37^ anti-correlation between LC activity levels and sigma (10-15Hz) power representing sleep spindle activity (**Fig. 4B**).

**Figure 4.**
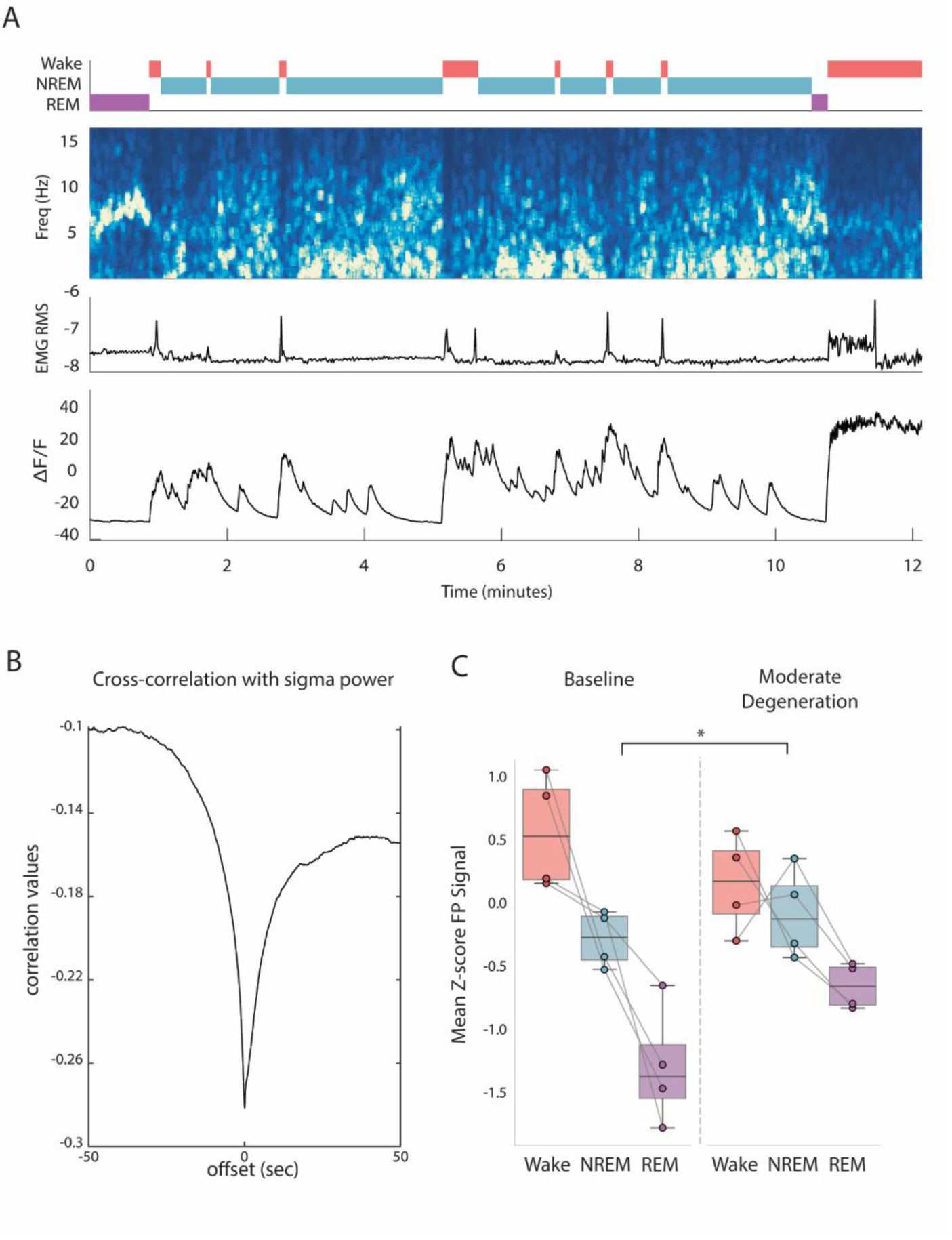
LC degeneration alters state-dependent LC photometry signal. (A) Representative sleep and LC dynamics across 12 minutes in one mouse at baseline. Top, hypnogram (time-course) of vigilance states. Red, wake; Cyan, NREM sleep; Purple, REM sleep. Second row shows corresponding EEG spectrogram (time-frequency decomposition). Third row: EMG RMS. Note micro-arousals during NREM sleep and elevated muscle activity during wakefulness. Fourth row: LC GCaMP population calcium activity. Note infraslow oscillations during NREM sleep, minimal levels in REM sleep, and high levels around arousals and wakefulness. (B) LC GCaMP cross-correlation with NREM sigma power. (C) Mean Z-scored LC signal (y-axis) by state (x-axis) at baseline (left) vs. moderate degeneration (right), n=4 mice; Difference between LC dynamicsin wake vs. REM sleep decreased significantly (p = 0.00309).

Comparing baseline to moderate degeneration, the difference between average LC GCaMP signal levels across wakefulness and REM sleep reduced significantly (p = 0.00309 via permutation test, Methods). This pattern (**Fig. 4C**) is consistent with a decrease in wakefulness and an increased activity in REM sleep, further supported by a significant Session × State interaction in a two-way repeated-measures ANOVA (p = 0.0088; Methods).

Together these findings show that despite the modest sample size, monitoring LC calcium dynamics reveals robust changes that occur upon moderate structural LC degeneration, occurring in parallel with REM sleep impairments.

## Discussion

To establish a causal, controlled model of gradual and partial LC degeneration, we expressed Cre-dependent Caspase-3 in LC neurons of female DBH-Cre mice. Diluting the viral dose produced a graded reduction in LC survival, and using our selected dose, we characterized a clear time-course of degeneration: negligible loss within the first three days, mild degeneration (∼25%) by day 8 post injection, and moderate degeneration (∼50%) by day 18. Applying this same time-course to a separate cohort implanted for sleep and LC photometry recordings, we found that progressive LC loss altered REM sleep features: REM sleep fragmented into shorter, more frequent episodes already at the mild stage. REM theta power declined progressively across time, and by the moderate stage, these impairments were accompanied by a modest increase in total REM sleep during the light period. The increase in total REM sleep may represent a compensatory response to impaired REM organization, although other explanations, including altered REM pressure or redistribution across shorter episodes, cannot be excluded.

Where do our results stand with respect to previous studies examining LC damage and sleep? Prior studies probing the LC’s role in sleep-wake regulation have used a range of chemical lesion methods with differing specificity, such as DSP4^21,25^, 6-OHDA^22,23^ and anti-DBH-saporin^38^; and have sampled sleep at time points ranging from 1–4 days to 3 months post-lesion. Even so, a common theme across these studies is that extensive, rapid LC damage has little lasting effect on sleep architecture beyond the first few days post-lesion^21,27,38–42^. Whereas sleep changes have been reported, they are inconsistent: some studies find increased, fragmented REM sleep during dark hours^26^ alongside decreased REM during light hours^25,38^, while others report the opposite pattern of increased light-hour REM^23^. Our findings are consistent with this broader pattern: LC degeneration did not produce a major disruption of overall sleep architecture. However, the light-hour REM fragmentation we found has, to our knowledge, not been previously reported. Unlike the acute, near-complete lesions used in prior studies, our model produces damage that is both gradual and partial — properties that may engage different circuit-level compensatory mechanisms than those triggered by rapid, extensive LC loss ^43–45^, and that may more closely approximate the time course and extent of LC degeneration in progressive neurodegenerative disease. A further methodological difference is that our study used polysomnography in mice, whereas most previous LC-lesion studies were conducted in rats; species and methodological differences may therefore also contribute to the discrepancy with prior work In the clinical neurodegeneration context, LC degeneration is an early feature of several neurodegenerative conditions, most extensively documented in Alzheimer’s disease (AD), where sleep is disrupted in several ways, including a general reduction in REM sleep^6^. Our results of REM fragmentation and, at the moderate stage, increased total REM sleep do not directly match this profile. However, theta power reduction in REM is consistent with reports in older adults linking increased tonic LC activity to reduced REM theta oscillations. Those studies did not directly assess LC structural integrity, but the convergence raises the possibility that the LC–theta relationship we observe here at the structural level parallels changes in LC activity in human aging. More broadly, REM fragmentation was among the most sensitive markers of LC degeneration in our model, suggesting finer-grained analysis of REM structure could be informative in patients. REM sleep episode length and its fragmentation are not typically tracked as systematic outcome measures in patient studies, and could constitute a sensitive readout beyond commonly used macro-sleep metrics.

What might drive the REM fragmentation? Because LC activity is closely associated with arousal and sleep-to-wake transitions^46^, increased noradrenergic activity in surviving LC neurons, and/or altered sensitivity of post-synaptic noradrenergic receptors, could reduce REM-state stability or increase susceptibility to premature arousal, thereby contributing to the shorter and more frequent REM episodes. Indeed, a similar compensatory mechanism has been proposed in humans, where surviving LC neurons are thought to undergo compensatory adaptations that increase noradrenergic signaling^47,48^. The reduced contrast between LC activity levels across wakefulness and REM sleep in our photometry recordings (**Fig. 4**) is compatible with this possibility. However, population-level fiber photometry cannot distinguish increased per-neuron activity from fewer neurons contributing to a population signal. Resolving this conclusively would require single-unit or genetically targeted approaches beyond the scope of this study.

Two limitations of the current study should be explicitly acknowledged. First, time elapsed over several weeks changed some aspects of sleep irrespective of the experimental manipulation (**Fig. S1C, E**). Accordingly, both control and Casp3-injected animals showed increased locomotor activity during dark hours across the recording sessions, likely reflecting a habituation process to the recording environment that completed over weeks. Future experiments using a design that allows recovery and habituation before the induction of degeneration (such as an inducible viral vector) can help resolve this issue. Second, this study used only female mice; given the known sex differences in sleep architecture and neurodegenerative disease trajectories, future studies are needed to determine whether these findings extend to male mice.

Together, these results show that gradual, partial LC degeneration is sufficient on its own to disrupt REM sleep architecture and attenuate REM theta oscillations. This isolated effect does not reproduce the broader sleep pathology seen in AD, such as insomnia and overall REM suppression, suggesting that LC loss alone is not the primary driver of the entire clinical phenotype, and that disease-associated sleep disturbance more likely depends on LC degeneration compounding with other pathological processes over time. Testing this directly will require complementary models; for example, applying similar longitudinal sleep and photometry approaches to transgenic AD models in which pathology targets other subcortical neuromodulatory nuclei while sparing the LC^19^ could help isolate each system’s individual contribution to the sleep phenotype of AD, and clarify whether their combined degeneration is needed to reproduce it in full. REM fragmentation and theta attenuation may therefore serve as tractable, LC-specific readouts for studying the functional consequences of LC degeneration, independent of any single disease model, and for testing candidate interventions at defined stages of cell loss.

## Acknowledgements

The work was supported by grants to Y.N. from the Israel Science Foundation (ISF 1557/22) and the European Research Council (ERC-2019-COG 864353). Additionally, the Nir lab is supported by the Sieratzki Institute for Advances in Neurosciences.

## Author Contributions

- Conception and design of research: N. Zelinger, N. Regev, and Y. Nir.
- Funding acquisition: Y. Nir.
- Data collection: N. Zelinger, U. Kimchy, N. Gabay
- Data analysis: N. Zelinger and E. Burian
- Manuscript Writing: N. Zelinger, N. Regev, and Y. Nir.
- Critical Review of Results and Manuscript Comments: All authors.

## Supplementary data

**Supplementary Table 1:** List of all animals in virus dilution verification cohort.

| ID | Injected hemisphere<br># cells | Uninjected<br>hemisphere # cells | titter | number<br>of slices | percent of<br>LC cells<br>surviving |
| --- | --- | --- | --- | --- | --- |
| 10 | 359 | 510 | x3 | 6 | 70.39 |
| 9 | 478 | 566 | x3 | 6 | 84.45 |
| 8 | 575 | 776 | x3 | 6 | 74.10 |
| 7 | 395 | 933 | x3 | 6 | 42.34 |
| 6 | 336 | 459 | x10 | 6 | 73.20 |
| 5 | 343 | 549 | x10 | 6 | 62.48 |
| 4 | 369 | 492 | x10 | 6 | 75.00 |
| 3 | 279 | 394 | x10 | 6 | 70.81 |
| 2 | 165 | 199 | x100 | 6 | 82.91 |
| 1 | 276 | 322 | x100 | 6 | 85.71 |

**Supplementary Table 2:** List of all animals used the timeline characterization cohort.

| ID | Injected hemisphere<br># cells | Non-injected<br>hemisphere # cells | days | number<br>of slices | percent of LC<br>cells surviving |
| --- | --- | --- | --- | --- | --- |
| 1 | 642 | 645 | 3 | 12 | 100.47 |
| 2 | 517 | 524 | 3 | 5 | 101.35 |
| 3 | 910 | 535 | 8 | 16 | 58.79 |
| 4 | 818 | 577 | 8 | 15 | 70.54 |
| 5 | 1183 | 857 | 8 | 15 | 72.44 |
| 6 | 912 | 950 | 8 | 18 | 72.44 |
| 7 | 700 | 554 | 8 | 9 | 79.14 |
| 8 | 998 | 810 | 8 | 21 | 81.16 |
| 9 | 544 | 482 | 8 | 10 | 88.60 |
| 10 | 874 | 212 | 18 | 16 | 24.26 |
| 11 | 1100 | 500 | 18 | 16 | 45.45 |
| 12 | 1834 | 952 | 18 | 16 | 51.91 |
| 13 | 2053 | 1074 | 18 | 20 | 52.31 |
| 14 | 678 | 422 | 18 | 8 | 62.24 |
| 15 | 1419 | 957 | 18 | 16 | 67.44 |
| 16 | 1876 | 1271 | 18 | 19 | 67.75 |

**Supplementary Figure 1.**
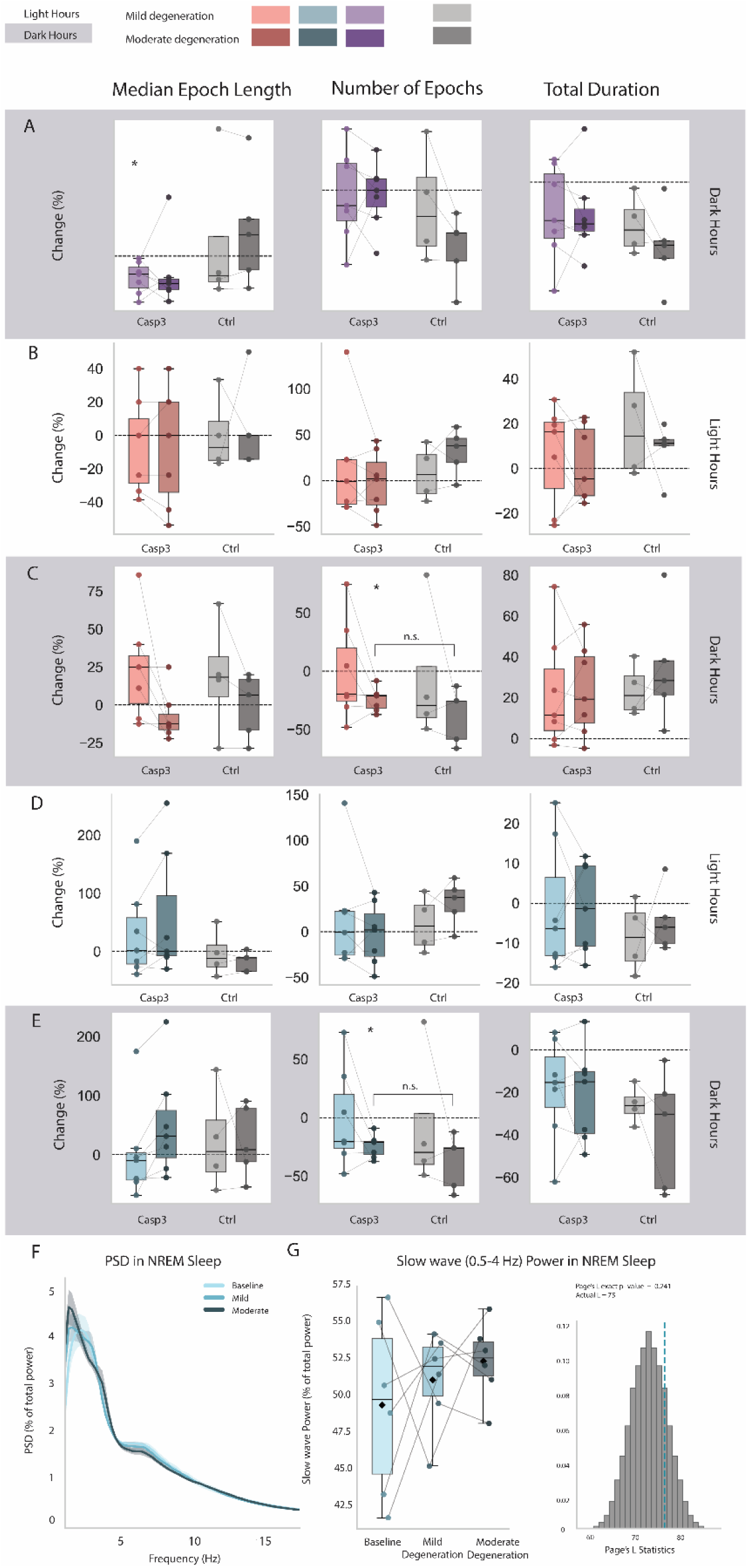
State- and metric-specific changes in sleep architecture across degeneration stages. Percent change from baseline in epoch metrics by state, hours, and time: (A) REM sleep in dark phase; (B) Wake in light phase; (C) Wake in dark phase; (D) NREM sleep in light phase; (E) NREM sleep in dark phase. Asterisks indicate significant within-group change from baseline; "n.s." indicates a non-significant between-group comparison to control. (F) EEG PSD during NREM sleep. (G) NREM slow-wave power (0.5–4 Hz) across degeneration stages, showing no significant trend (Page’s L trend test, L = 75, n = 6, p = 0.24). Data are mean ± SEM

